# Adaptive evolution of anti-viral siRNAi genes in bumblebees

**DOI:** 10.1101/017681

**Authors:** Sophie Helbing, H. Michael G. Lattorff

## Abstract

The high density of frequently interacting and closely related individuals in social insects enhance pathogen transmission and establishment within colonies. Group-mediated behavior supporting immune defenses tend to decrease selection acting on immune genes. Along with low effective population sizes this will result in relaxed constraint and rapid evolution of genes of the immune system. Here we show that sociality is the main driver of selection in antiviral siRNAi genes in social bumblebees compared to their socially parasitic cuckoo bumblebees that lack a worker caste. RNAi genes show frequent positive selection at the codon level additionally supported by the occurrence of parallel evolution and their evolutionary rate is linked to their pathway specific position with genes directly interacting with viruses showing the highest rates of molecular evolution. We suggest that indeed higher pathogen load in social insects drive adaptive evolution of immune genes, if not compensated by behavior.

## Introduction

Social insects, like ants, bees, wasps and termites, represent an extremely successful group. Although they represent only 2 % of all insect (invertebrate) species, they contribute up to 25% of the total insect biomass (Wilson 1990). This success is mainly attributed to the division of labor amongst workers, but also to the reproductive division of labor between castes with queens monopolizing reproduction whereas workers remain functionally sterile (Lattorff and Moritz 2013). This reproductive division of labor is enhanced by the high degree of relatedness allowing kin selection (Hamilton 1964) to work efficiently.

However, this system might represent also some major drawbacks. Social insects are a prime target for parasites (Schmid-Hempel 1998), as their transmission is enhanced due to frequent social interactions and their establishment might be augmented by the high relatedness within colonies as well as by the high degree of nest homeostasis (Boomsma et al. 2005). Despite the high parasite pressure, a lack of immune system genes has been recognized within all the completely sequenced social insect genomes when compared to other, non-social insect species (Evans et al. 2006; Smith et al. 2011). Several explanations have been put forward in order to explain this discrepancy. The most prominent one explains the lack of immune genes by an advanced level of socially mediated physiological adaptations and behavioral activities directed against intruding parasites, summarized as “social immunity” (Cremer et al. 2007). Indeed, a huge range of activities has been described that altogether might reduce the selection coefficient acting on genes of the innate immune system. In combination with drastically lowered effective population sizes in social insects (Romiguier et al. 2014) due to the reproductive division of labor, this might result in reduced or even absent purifying selection on immune genes (Evans et al. 2006). This will result in non-efficient removal of slightly deleterious mutations (also known as relaxed constraint) that will behave like nearly neutral mutations (Ohta 1987) so that non-synonymous substitutions will increase in frequency and finally contribute to non-synonymous divergence between species. The latter one has been frequently observed for social insects implying faster evolution of immune genes (Harpur & Zayed 2013), especially when compared to solitary insects like *Drosophila* (Viljakainen et al. 2009).

Nevertheless, comparisons between highly eusocial ants or the honeybee (*Apis mellifera*) with solitary insects like *Drosophila* or *Anopheles* suffer from several confounding factors. Hymenopteran social insects are haplo-diploid (females are diploid whereas males are haploid), a genetic constitution that by itself reduces the effective population size (Crozier 1977). Furthermore, these lineages might have had a common ancestor 300 million years ago and might also show lineage specific effects.

We study the effect of sociality on the rates of evolution in a superior system consisting of social bumblebees and their socially parasitic cuckoo bumblebees, the latter one are non-social due to the lack of a worker caste. The eusocial lineages are characterized by annual colonies headed by one single-mated queen. The colony cycle is composed of different stages: (I.) nest establishment; (II.) production of a sterile worker caste ensuring colony growth via division of labor and (III.) towards the end of the season production of sexual offspring (Goulson 2010). The cuckoo bumblebee females enter host nests at a time point during early development of the social nest and kill the resident queen in order to lay their own eggs that will be cared for by the resident worker force. As this relationship is very specific with every cuckoo bumblebee species parasitizing a unique host species or at least only a restricted number of host species, a certain couple of host/parasite species do show similar life history characters and use a shared environment, at least for the time of shared nests. Cuckoo bumblebees will show an even more reduced effective population size than their hosts, as they never may parasitize all available nests (Erler and Lattorff 2010). This might result in even stronger non-synonymous divergence under relaxed constraint than it might be observed for social species. In a large comparison of several host/social parasite couples spread throughout the order of insects, Bromham and Leys (2005) have shown that indeed social parasites show higher rates of molecular evolution, at least in seven out of eight couples of species.

We are studying the rates of molecular evolution in genes of the anti-viral siRNAi pathway that is responsible for the degradation of double stranded RNA usually derived from viruses. This system is not inducible, as this pathway also interferes with the regulation of gene expression of host genes via microRNAs. This system, when acting in different castes like queens and workers, which might show antagonistic patterns of gene regulation, might additionally slow down the rates of evolution in these genes. Nevertheless, if parasites, in this case viruses, which are known to be widespread in social insects, in which interspecific transmission might also occur, impose a high selective pressure on them, then this might become visible by higher rates of evolution within the siRNAi genes in those species that are social.

### Materials and Methods

Six genes of the siRNAi pathway were partially sequenced and compared across six social *Bombus* species and six of their respective cuckoo bumblebees (*Bombus*, subgenus *Psithyrus*) with each species represented by a maximum of four haploid drones. Detailed information concerning the studied species, their geographic origin and method for DNA extraction has been published elsewhere (Erler et al. 2014).

In order to estimate intra-specific polymorphism 20 and 19 haploid males of *B. terrestris* and *B. vestalis*, respectively, as well as 5 diploid females of *B. lapidarius* (workers) and *B. rupestris* (females) were sequenced for one of the siRNAi genes (*r2d2*). All those individuals were unrelated to each other as revealed by microsatellite genotyping (data not shown).

#### Homology searches & Primer design

The RNAi genes were identified in the draft assembly of the *Bombus terrestris* genome (assembly: Bter 1.0, AELG01000001:AELG01010672) and the *Bombus impatiens* genome (assembly: Bimp 1.0, AEQM01000001-AEQM01091524) (International Bumblebee Genome Sequencing Consortium, 2014) using homology searches based on *Drosophila melanogaster* protein sequences and *Apis mellifera* DNA sequences (Honey Bee Genome, Assembly 4). Primers were designed on the consensus sequence resulting from the alignment of the *B. terrestris* and *B. impatiens* sequences using Primer3 (Rozen and Skaletsky 2000). Primers used in this study are reported in table S2.

#### PCR

PCR–protocols were chosen according to the fragment length of the target gene. Fragments with expected sizes above 2000 bp were amplified with long-range PCR utilizing TaKaRaLA Taq Polymerase (MoBiTec, Göttingen, Germany). Smaller fragments were amplified using PeqLab Gold Taq (Peqlab, Erlangen, Germany). PCR protocols are given in table S3.

#### Sequencing

Sequencing of *r2d2* (interspecific variation, 12 species), *vig* and *maelstrom* (*mael*) was performed by LGC Genomics (Berlin, Germany) using traditional Sanger sequencing. Samples were sequenced directly and in both directions. Sequence chromatograms were inspected and confirmed manually. Forward and reverse sequences were assembled using ContigExpress implemented in Vector NTI Advance 10.2.0 (Invitrogen, Karlsruhe, Germany) in order to detect PCR or sequencing errors.

PCR products of *argonaute2* (*ago2*), *armitage* (*armi*) and *dicer 2* (*dcr2*) were sequenced using the Ion Torrent PGM (Life Technologies). Therefore each species was tagged with a species-specific barcode adapter and for each gene all individuals per species were pooled in equimolar amounts. Finally, all amplicons per species were pooled together in equimolar amounts. The 100 base-read barcoded libraries were prepared using the Ion Xpress Barcode Adapters 1-16 Kit following the instructions of the Ion Xpress Plus gDNA and Amplicon Library Preparation user guide. CLC Bio Genomics Workbench 5.0 (CLC Bio, Aarhus, Denmark) was used to (I) map the reads to the *Bombus terrestris* reference sequences using default settings and (II) to detect SNP’s in the high-throughput sequencing data using the SNP detection tool. The latter was performed using default settings except for the minimum coverage and minimum variant frequency which were adjusted to the library (species /gene) specific requirements.

Sequencing of *r2d2* (intraspecific variation, 4 species) was done using the Ion Torrent PGM (Life Technologies). All amplicons within a species were pooled in equimolar amounts and species were barcoded using the Ion Xpress Barcode Adapters 1-16 Kit according to the instructions of the Ion Xpress Plus gDNA and Amplicon Library Preparation user guidelines. Pooled sequences were mapped to reference sequences for the respective species derived from Sanger sequencing. SNPs were detected using the Quality-based variant detection function implemented in CLC Genomics Workbench v7.0.4 (CLC Bio, Aarhus, Denmark) using the default settings except for the minimum frequency of variants, which was adjusted to the number of chromosomes and the average coverage.

For all species, except *B. perezi*, Cameron et al. 2007 provided sequences of four nuclear genes (*EF-1 alpha*, *Arginine kinase*-AK2, *rhodopsin*, *PEPCK*) on GenBank. Erler et al. 2014 provided *B. perezi* sequences (*EF-1 alpha*, *Arginine kinase*-AK2, *rhodopsin*, *PEPCK*, GenBank accession numbers: KC662163-67). Based on these genes the phylogenetic relationship between the focal species was reconstructed and they were also used to estimate the rate of evolution on non-immune genes.

#### Evolutionary analyses

Multiple sequence alignment was performed using ClustalW implemented in Mega 5.05 (Tamura et al. 2011). Rates of molecular evolution of the RNAi genes were calculated based on coding sequences separately for each gene.

Within species polymorphism as well as the ratio of the number of non-synonymous substitutions per non-synonymous sites to the number of synonymous substitutions per synonymous sites including Jukes-Cantor correction (Jukes and Cantor 1969) was estimated using DnaSP v5 (Librado and Rozas 2009). In order to compare the evolutionary rates between social and non social cuckoo bumblebees, sequences were furthermore classified according to their affiliation to either the social or parasite dataset. Stability analyses by means of jack-knifing over species were performed to confirm the values for each group.

#### Site-specific selection

Codeml, implemented in the PAML package (Yang 2007) providing models allowing *ω* to vary among sites, was used in order to identify sites under positive selection. Therefore, non-immune gene based phylogeny reconstruction was performed using PHYLIP version 3.69 (Neighbor-Joining method) (Felsenstein 2005). The comparisons comprised models M7 (beta) vs. M8 (beta &*ω*) and were repeated twice changing the initial *ω* (*ω*< 1; *ω*>1). Significance was assessed using a likelihood ratio test (LRT) comparing M7 and M8 models. We used the Bayes empirical Bayes (BEB) approach (codeml implementation) for the inference of site-specific selection based on calculated posterior probabilities of *ω* classes for each site.

#### Parallel evolution

The detection of signs of parallel evolution may also indicate the occurrence of positive selection. The reconstruction of the ancestral amino acid sequence, based on the phylogeny and the amino acid sequences of the present-day species enables the inference of the evolutionary pathway of amino acid substitutions at each site (Yang et al. 1995). Therefore, nucleotide sequences were translated and resulting amino acid sequences were aligned using ClustalW (Thompson et al. 1994) implemented in BioEdit (Hall 1999). The ancestral character reconstruction using the maximum likelihood (ML) algorithm (substitution model: Jones-Taylor-Thornton model, JTT), as well as the construction of the phylogenetic tree (Fig.4), based on the non-immune genes, was conducted with Mega 5.05 (Tamura et al. 2011). Hence, it was possible to infer changes that occurred on each lineage and therefore enabled the identification of shared amino acid replacements. In order to distinguish whether the observed change is attributable to chance alone or to parallel evolution, the program CAPE (Zhang and Kumar 1997) was used. CAPE computes the probability that the observed parallel substitutions are attributable to random chance alone - based on the amino acid sequences and the divergence between species according to a reference phylogenetic tree (Zhang and Kumar 1997). The CAPE computations were performed using the JTT substitution model. The required tree topology (PHYLIP format) as well as the notation of the nodes was verified using the program package ANCESTOR (Zhang and Nei 1997).

**Figure 4.**
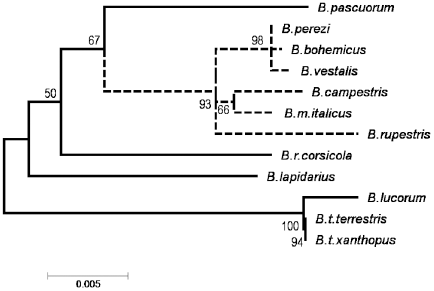
Bombus phylogenetics. Phylogenetic relationships of the six host (solid line) and social parasite (dashed line) bumblebee species based on the coding sequences of non-immune genes (Arginine kinase, EF-1 alpha, PEPCK, Rhodopsin) inferred using the Maximum Likelihood method based on the Kimura 2-parameter model method (including test of phylogeny: bootstrap with 500 replicates).

### Results

Six RNAi loci were partially amplified: *argonaute2* (*ago2*), *armitage* (*armi*), *dicer2* (*dcr2*), *maelstrom* (*mael*), *r2d2*, *vasa intronic gene* (*vig*), and coding sequences were compared across different *Bombus* species. Length of coding sequences analyzed as well as polymorphism data are reported in table S1. In summary, the total number of within-species polymorphism was markedly low and affect almost exclusively host species. For *B. t. xanthopus* we observed two synonymous substitutions in the coding region of *vig*; furthermore one non-synonymous (*ago2*) and one synonymous substitution (*r2d2*) in *B. lucorum*; always one synonymous polymorphism appeared in *B. terrestris* (*mael*) and *B. lapidarius* (*dcr2*) and one non-synonymous substitution in *B. bohemicus* (*r2d2*). In addition, InDel events appeared in the coding sequences of *armi* and *r2d2* (Table S1). Due to the low sample size representing each species this study fails to infer patterns of selection also considering population data. Nevertheless, at least we can draw on intra-specific polymorphism data for *r2d2* - sequenced in 4 *Bombus* species: *B. terrestris* (drones, N = 20); *B.vestalis* (drones, N = 19); *B. lapidarius* (workers, N = 5) and *B. rupestris* (workers, N = 5). These data confirm the observed lack of diversity within coding regions.

#### Evolutionary rates of anti-viral siRNAi genes

In order to detect footprints of selection in siRNAi genes, the overall ratio of non-synonymous- to synonymous substitutions was calculated and compared with the evolutionary rate of non-immune genes (*Arginine kinase*, *EF-1 alpha*, *PEPCK* and *Rhodopsin*;). The studied siRNAi genes showed an elevated rate of molecular evolution compared to the non-immune genes (mean_RNAi_ K_A_/K_S_ = 0.35; mean_non-imm_ K_A_/K_S_ = 0.02; Mann-Whitney-U-Test: N = 10, Z = 2.56; p = 0.01). We examined if higher values of this ratio are driven by non-synonymous changes and indeed, K_A_ is 28 times higher in RNAi genes compared to non-immune genes (t-test; K_A_: mean_RNAi_=0.028; mean_non-imm_= 0.001; *P* < 0.05), while Ks is not significantly different (t-test; K_S_: mean_RNAi_= 0.101; mean_non-imm_= 0.081; *P* > 0.05). Additionally, non-immune genes showed consistently low K_A_/K_S_ values (0.01-0.03) indicating strong purifying selection, while the rate of adaptive evolution in RNAi genes varied. The strongest signs of adaptive evolution were observed for *r2d2* (K_A_/K_S_ = 0.52) and *ago2* (K_A_/K_S_ = 0.65), both encoding proteins directly interacting with viral components and are therefore expected to be hotspots of adaptive changes. A slightly elevated rate of adaptive evolution was observed for *dcr2* (K_A_/K_S_ = 0.29) and the helicase *armi* (K_A_/K_S_ = 0.43). In contrast, we also identified genes (*vig*, *mael*) showing the occurrence of purifying selection indicated by low K_A_/K_S_ values. Especially *vig,* involved in the RNAi defense mechanism but also in heterochromatin formation (Caudy et al.2002; Gracheva et al. 2009), showed a low K_A_/K_S_ value compared to the non-immune genes. Besides that it is the smallest of all genes analyzed reducing the likelihood to pick up a sufficient number of changes and it is located within the intron of another gene (*vasa*) exposing it to selective pressures acting on this gene.

#### Rates of molecular evolution as a function of life-history

Reduced selective constraint is expected to increase non-synonymous divergence in socially parasitic species. Astonishingly, we observed faster evolution for three out of six genes for the social species, while only two siRNAi genes showed higher rates of molecular evolution in the non-social parasite species (Fig. 1). Thus, social host species showed an elevated rate of adaptive evolution relative to the socially parasitic cuckoo bumblebees for *ago2* (K_A_/K_S_ (Host): 0.91; K_A_/K_S_ (Parasites): 0.39; T-Test, P = 9 × 10^-6^), *mael* (K_A_/K_S_ (Host): 0.34; K_A_/K_S_ (Parasites): 0.03; T-Test, P = 2 × 10^-6^) and *armi* (K_A_/K_S_ (Host): 0.45; K_A_/K_S_ (Parasites): 0.16; T-Test, P = 1.84 × 10^-4^), while for *dcr2* (K_A_/K_S_ (Host): 0.27; K_A_/K_S_ (Parasites): 0.74; T-Test, P = 1.44 × 10^-3^) and *vig* (K_A_/K_S_ (Host): 0.07; K_A_/K_S_ (Parasites): 0.16; T-Test, P = 1 × 10^-6^) parasites’ K_A_/K_S_ ratios significantly exceed the values for social species. Comparing the strength of selection between social hosts and parasites for *r2d2* showed that the evolutionary rate is not significantly different (K_A_/K_S_ (Host): 0.54; K_A_/K_S_ (Parasites): 0.48; T-Test, P = 0.08) indicating high selective pressures supported by the complete lack of within species polymorphism (see above).

**Figure 1.**
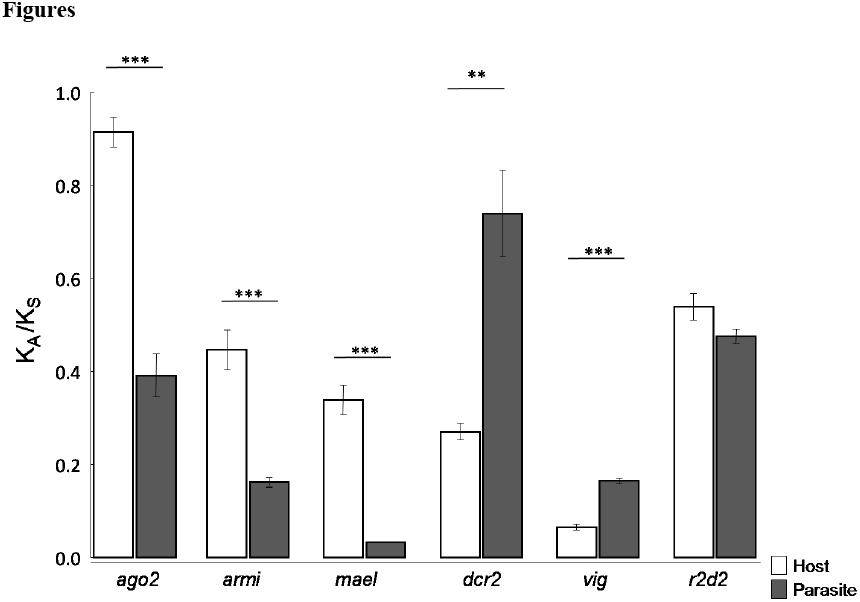
Evolutionary rate of siRNAi genes for social and socially parasitic cuckoo bumblebees. Ka/Ks values by means of jack-knifing over species for the social (open bars) and non-social (filled bars) datasets. Statistically (T-Test) supported differences: ** P-value<0.01; *** P-value <0.0001.

#### Codon-based positive selection & parallel evolution

None of the studied RNAi genes had (global) K_A_/K_S_ > 1 – indicative for positive selection. Nevertheless, we found evidence for positively selected sites within the coding region – especially for *r2d2* and *ago2*. In the ML-based approach we compared model M7 vs. M8 in order to detect sites experiencing positive selection. For *ago2*, *r2d2*, *dcr2* and *armi* the LRTs suggests a significant difference between the compared models, indicating the presence of sites evolving under positive selection. As parallel evolving sites might also serve as a hint for positive selection, substitution events shared between the social species and their social parasites were determined. Our data do not support any general pattern of parallel evolution, neither between cuckoo bumblebees and their respective hosts nor with respect to geographical origin. Nevertheless, the occurrence of parallel substitutions was observed for each locus, except for *vig* (Table 1, Table 2).

**Table 1:**
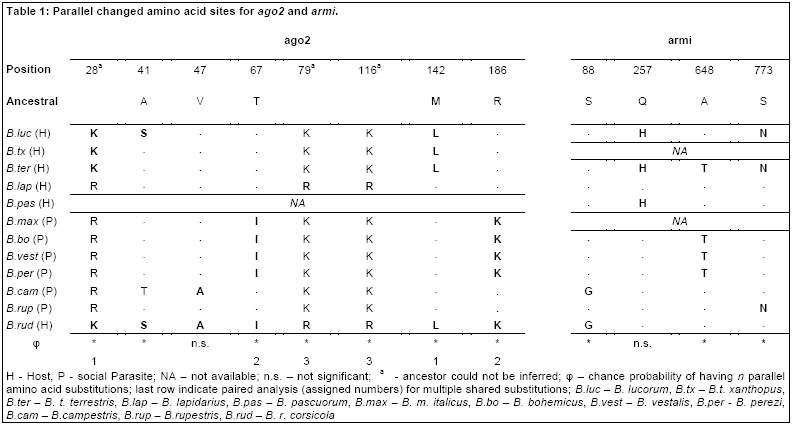
Parallel changed amino acid sites for *ago2* and *armi*.

|  | **ago2** |  |  |  |  |  |  |  | **armi** |  |  |  |
| --- | --- | --- | --- | --- | --- | --- | --- | --- | --- | --- | --- | --- |
| **Position** | 28<sup>a</sup> | 41 | 47 | 67 | 79<sup>a</sup> | 116<sup>a</sup> | 142 | 186 | 88 | 257 | 648 | 773 |
| **Ancestral** |  | A | V | T |  |  | M | R | S | Q | A | S |
| *B.luc* (H) | **K** | **S** | . | . | K | K | **L** | . | . | **H** | . | **N** |
| *B.tx* (H) | **K** | . | . | . | K | K | **L** | . | NA |  |  |  |
| *B.ter* (H) | **K** | . | . | . | K | K | **L** | . | . | **H** | **T** | **N** |
| *B.lap* (H) | R | . | . | . | **R** | **R** | . | . | . | . | . | . |
| *B.pas* (H) | NA |  |  |  |  |  |  |  | . | **H** | . | . |
| *B.max* (P) | R | . | . | **I** | K | K | . | **K** | NA |  |  |  |
| *B.bo* (P) | R | . | . | **I** | K | K | . | **K** | . | . | **T** | . |
| *B.vest* (P) | R | . | . | **I** | K | K | . | **K** | . | . | **T** | . |
| *B.per* (P) | R | . | . | **I** | K | K | . | **K** | . | . | **T** | . |
| *B.cam* (P) | R | T | **A** | . | K | K | . | . | **G** | . | . | . |
| *B.rup* (P) | R | . | . | . | K | K | . | . | . | . | . | **N** |
| *B.rud* (H) | **K** | **S** | **A** | **I** | **R** | **R** | **L** | **K** | **G** | . | . | . |
| $\phi$ | * | * | n.s. | * | * | * | * | * | * | n.s. | * | * |
|  | 1 |  |  | 2 | 3 | 3 | 1 | 2 |  |  |  |  |

H - Host, P - social Parasite; NA – not available; n.s. – not significant; <sup>a</sup> - ancestor could not be inferred; $\phi$ – chance probability of having *n* parallel amino acid substitutions; last row indicate paired analysis (assigned numbers) for multiple shared substitutions; *B.luc* – *B. lucorum*, *B.tx* – *B.t. xanthopus*, *B.ter* – *B. t. terrestris*, *B.lap* – *B. lapidarius*, *B.pas* – *B. pascuorum*, *B.max* – *B. m. italicus*, *B.bo* – *B. bohemicus*, *B.vest* – *B. vestalis*, *B.per* - *B. perezi*, *B.cam* – *B.campestris*, *B.rup* – *B.rupestris*, *B.rud* – *B. r. corsicola*

**Table 2:** Parallel changed amino acid sites for *dcr2*, *mael* and *r2d2*.

|  | *dcr2* |  |  | *mael* | *r2d2* |  |  |  |  |  |  |
| --- | --- | --- | --- | --- | --- | --- | --- | --- | --- | --- | --- |
| Position | 56 | 79<sup>a</sup> | 82<sup>a</sup> | 29 | 54 | 103 | 87<sup>a</sup> | 119<sup>a</sup> | 134<sup>a</sup> | 149<sup>a</sup> | 161<sup>a</sup> |
| Ancestral | R |  |  | N | T | Y |  |  |  |  |  |
| *B.luc* (H) | **K** | N | H | . | . | . | **N** | T | L | **T** | **S** |
| *B.tx* (H) | **K** | N | H | . | **I** | **H** | **N** | T | L | **T** | **S** |
| *B.ter* (H) | NA |  |  | . | . | **H** | **N** | T | L | **T** | **S** |
| *B.lap* (H) | . | N | L | **D** | S | . | S | T | I | **T** | N |
| *B.pas* (H) | **K** | Y | L | **S** | **I** | **H** | G | **I** | L | I | N |
| *B.max* (P) | . | Y | F | . | . | . | T | T | I | I | N |
| *B.bo* (P) | . | Y | F | . | . | . | T | T | I | I | N |
| *B.vest* (P) | . | Y | F | . | . | . | T | T | I | I | N |
| *B.per* (P) | . | Y | F | . | . | . | T | T | I | I | N |
| *B.cam* (P) | . | Y | F | **D** | . | . | T | T | I | I | N |
| *B.rup* (P) | . | **C** | **S** | **D** | . | . | **N** | T | I | I | **S** |
| *B.rud* (H) | . | **C** | **S** | **S** | . | . | T | **I** | L | **T** | N |
| φ | n.s. | * | * | *, * | * | * | * | n.s. | n.s. | n.s. | * |
|  |  | 4 | 4 | 5,6 |  |  | 7 |  |  |  | 7 |

H - Host, P - social Parasite; NA – not available; n.s. – not significant; <sup>a</sup> - ancestor could not be inferred; φ – chance probability of having *n* parallel amino acid substitutions; last row indicate paired analysis (assigned numbers) for multiple shared substitutions; *B.luc* – *B. lucorum*, *B.tx* – *B.t. xanthopus*, *B.ter* – *B. t. terrestris*, *B.lap* – *B. lapidarius*, *B.pas* – *B. pascuorum*, *B.max* – *B. m. italicus*, *B.bo* – *B. bohemicus*, *B.vest* – *B. vestalis*, *B.per* – *B. perezi*, *B.cam* – *B. campestris*, *B.rup* – *B. rupestris*, *B.rud* – *B. r.corsicola*

The comparison between models M7 and M8 for *ago2* yielded a LRT statistic of 16.76, significant at p = 0.001, with a proportion of sites (6.3%) with ω = 7.08 (Table S4). The ML analysis for *ago2* revealed three positively selected sites with *P*> 95% (BEB: sites 41 and 68 with *P*> 95 %; site 77 with *P*> 99%). All these sites are located in the nucleic acid binding domain PAZ (Fig. 2). Furthermore, five parallel changed sites were detected, always in pairs including *B. ruderatus*. Interestingly, two identical substitutions (sites 67 and 186) were shared between *B. ruderatus* and most of the social parasites except for *B. campestris* and *B. rupestris,* with one of them (site 67) showing an alteration of the chemical property of the amino acid (Table 1).

**Figure 2.**
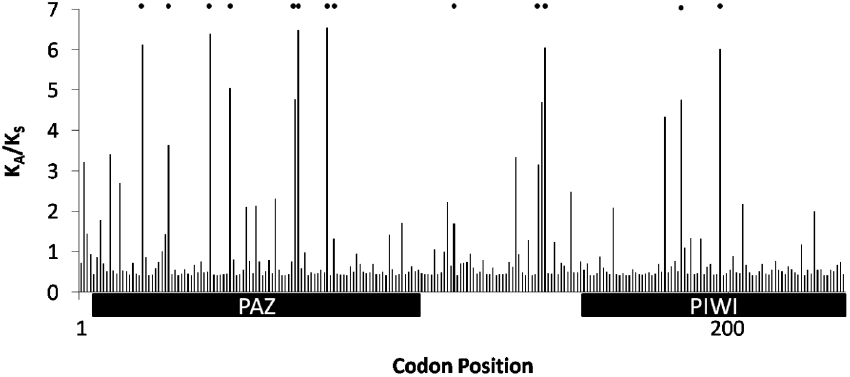
Evidence for positively selected sites in *ago2*. Posterior mean *ω* for each codon position of *ago2* under model M8. Symbols refer to positively selected sites (*ω* > 1) with posterior probabilities under model M8 (BEB method): *P* > 0.75 (BEB; sites: 20, 47, 144, 198); P > 0.95 (BEB; sites: 41, 68, 77) and parallel changed sites: 28^a^, 41, 67, 79^a^, 116^a^, 142, 186. ^a^ - ancestor could not be inferred. Bars below the plot indicate conserved domains.

The comparison of M7 vs. M8 for *r2d2* yielded a test statistic of 9.86, significant at p = 0.01 (Table S4). Here, M8 identified 30 positive selected sites (*P* > 50%) – three of them with *P* > 95% (BEB: site 87, prob. > 99%; sites 90 and 181, prob. > 95%; Fig. 3). Seven identical changes were observed, but only four of them passed the test for parallel evolution. One charge-altering substitution, which takes place in the *dsrm* domain, occurred in *B. t. xanthopus* and *B. pascuorum* (T54I, Ф= 0.014). *B. terrestris*, *B. t. xanthopus* and *B. pascuorum* shared the same charge altering amino acid replacement at position 103 (Y103H), unaccountable to chance alone, (Ф= 0.002) and thus strongly suggesting the occurrence of parallel evolution. Furthermore, there are two parallel-change sites (T87N; N161S; Ф= 0.000) in the *B. rupestris*-*B. lucorum/B. terrestris/B. t. xanthopus* comparison. However, it was not possible to infer the ancestral state at these positions.

**Figure 3.**
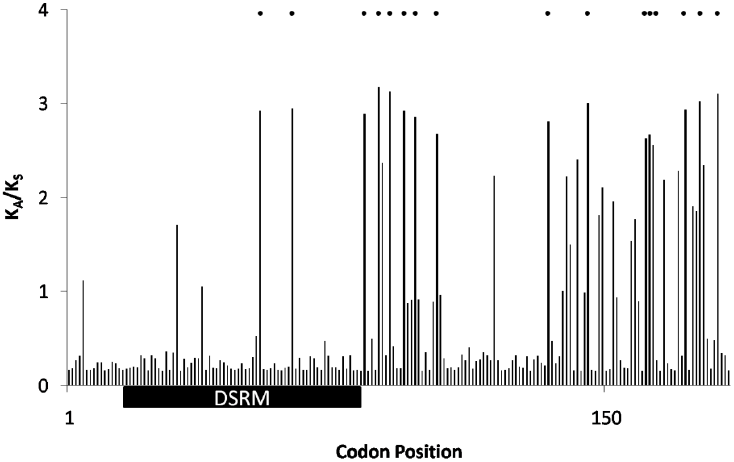
Evidence for positively selected sites in *r2d2*. Posterior mean *ω* for each codon position of *r2d2* (model M8). Symbols refer to positively selected sites (*ω* > 1) with posterior probabilities under model M8 (BEB method): *P* > 0.75 (BEB; sites: 54, 63, 83, 94, 97, 103, 134, 145, 161, 162, 163, 172, 176); P > 0.95 (BEB; sites: 87, 90,181) and parallel changed sites: 54, 87^a^, 103, 119^a^, 134^a^, 149^a^, 161^a^; ^a^ - ancestor could not be inferred. Bars below the plot indicate conserved domains.

For *dcr2* (M7 vs. M8: LRT= 15.38, significant at p = 0.001; Table S4) the BEB identified two sites (79, 82) to be positively selected with a probability of > 99%. Both sites are exceptionally variable and we observed potential parallel substitutions. Unfortunately, it was not possible to infer the ancestral state for both sites. Nevertheless, we calculated the probability for observing two identical substitutions shared between *B. ruderatus* and *B. rupestris* – as both species even showed identical substitutions at both sites. Testing for parallel evolution revealed that these changes are highly significant exceeding random chance expectation.

## Discussion

Life-history traits associated with sociality, infection risk within groups, are predicted to affect the rates of molecular evolution. Several studies (Viljakainen et al. 2009; Bulmer and Crozier 2004, 2006; Harpur and Zayed 2013; Erler et al. 2014) targeted at quantifying host parasite conflict-effects on the evolution of genes involved in antimicrobial or antifungal defense mechanisms in social insects. However, antiviral defense mechanisms (e.g. RNAi) are also part of the innate immune system leading to the inhibition of expression of viral genes essential for viral replication. Counteracting this, viruses encode essential virulence factors, called viral suppressors of RNAi (VSR) interfering with key-steps of host immune responses (Li and Ding 2006). Virus-associated characteristics (high replication and mutation rates) as well as VSR impose severe requirements on the host immune-system resulting in an evolutionary arms-race between components of the RNAi-pathway and viruses (Moissiard et al. 2004; Li et al. 2005; Obbard et al. 2006; Obbard et al. 2009a). For *Drosophila*, high rates of adaptive evolution of siRNAi genes are indicative for these co-evolutionary interactions (Obbard et al. 2006; Obbard et al. 2009b, Kolaczkowski et al. 2011).

We studied the evolutionary pattern of antiviral RNAi genes across the genus *Bombus*, as it comprises social-as well as non-social lineages (cuckoo bumblebees). A global comparison of K_A_/K_S_ values between siRNAi and non-immune genes revealed that the former evolve significantly faster indicating permanent exposure to selection pressure imposed by viruses, especially as insect infecting (RNA) viruses are emerging as serious threats (Singh et al. 2010; Levitt et al. 2013; Fürst et al. 2014). Antiviral RNAi mechanisms are activated in the presence of viral double-stranded (ds)-RNA, leading to the generation of small interfering RNAs (siRNAs) serving as template for RISC-mediated viral mRNA degradation (Meister and Tuschl 2004; Buchon and Vaury 2006; Ding 2010; Nayak et al. 2013). The studied siRNAi genes showed huge variance in their evolutionary rates, possibly reflecting their different functions. In that regard we identified genes potentially under purifying selection, especially *vig* and *mael*, while others showed elevated rates of evolution, even though none of them with K_A_/K_S_> 1 for the whole gene. The highest rates of adaptive evolution were detected for *r2d2* and *ago2*. *R2d2* functions in siRNA production, while *ago2* (core element of the RISC) mediates viral mRNA degradation. Thus, both gene products are required for an effective RNAi and represent potential targets for VSR (Nayak et al. 2010; van Mierlo et al. 2012).

The overall trend for the rate of adaptive evolution is largely consistent with results reported for *Drosophila* (Obbard et al. 2009). Nevertheless, *mael* was assigned to five per cent of the fastest evolving genes, while the rate of adaptive evolution in bumblebees is just slightly elevated compared to non-immune genes.

Although, we could not detect signals of positive selection for full genes, we found candidates with positively selected sites. For *r2d2* we identified 16 positively selected sites (BEB; *P*> 75%), interestingly only three fall within the *DSRM* domain. This is consistent with the outcome of a study by Kolaczkowski et al. (2011), who found a large proportion of adaptive changes in the region separating both *dsrm* domains, while just a few adaptive protein-coding changes in the N-terminal *dsrm* domain were detected. The relative abundance of adaptive changes in the *dsrm* domain might be driven by the removal of deleterious mutations, since point mutations within this domain abolished dsRNA binding (Liu et al. 2003). By contrast, the *ago2* PAZ domain showed an elevated proportion of positively selected sites. This leads to the suggestion that this rapidly changing domain might be a target of parasite-immune evasion strategies, especially considering that *ago2* is a target of viral suppressors of RNAi (VSR) as shown for *Drosophila* (Nayak et al. 2010; Barribeau et al. 2014). Nevertheless, sites under positive selection, identified in a comparison of highly (*Apis mellifera, A. florea*) and primitively eusocial (*B. terrestris, B. impatiens*) as well as a non-social (*Megachile rotundata*) species, either on the branch leading to sociality (social vs. non-social species) or on the branch to bumblebees (*Bombus* vs. *Apis*) (Barribeau et al. 2014) were not recovered as being positively selected in our study suggesting that these sites have been spread to fixation before the diversification within the genus *Bombus*.

Based on stability analyses, one species – *B. rupestris* - was identified to have a drastic impact on the estimated values of the social parasite species for *ago2* and *dcr2*: Excluding *B. rupestris* in case of *ago2* will lead to an increase of the parasite diversity, while in case of *dcr2* this diversity is reduced.

Furthermore, five out of six genes showed evidence for parallel changed substitutions across the genus *Bombus* - also indicative for positive selection. Parallel substitutions were observed within host species belonging to different clades as well as between social parasites and clades of the social host. In some cases shared substitutions affect multiple sites in a gene; for instance we observed two substitutions in close proximity in *dcr2* shared between *B. rupestris* and *B. ruderatus* (Table 2). We hypothesized positive selection might favor same sites in cuckoo bumblebees and their respective hosts, as they show similarities in their life-cycles, share the same environment and are therefore predicted to be exposed to similar pathogens. We also expected to identify parallel changed sites for species originating from the same sampling site, reflecting geographically-restricted selective pressure due to the occurrence of location-specific viruses. However, no evidence was found for parallel substitution patterns reflecting either the relationship between host-social parasite couples, or location-restricted adaptation.

Sociality is expected to influence the molecular evolution of genes in a number of ways: mainly through reduction in N_e_ due to reproductive monopolization, but also continuous oogenesis and/or caste biased gene expression. Utilizing a comparative approach encompassing social and non-social lineages throughout the order of insects (including host/social parasite couples), Bromham and Leys (2005) investigated sociality-associated effects on the rate of molecular evolution. Here, evidence for an effect of lower effective population size is rising, as eusocial Hymenoptera showed increased evolutionary rates compared to their nonsocial relatives, but decreased rates compared to social parasites experiencing a further reduction in N_e_ (Bromham and Leys 2005). However, for the studied anti-viral siRNAi genes the impact of low N_e_ is not that conclusive: for three out of six genes the rate of molecular evolution is considerably faster in social species, even though assumed to have a higher N_e_ (Erler and Lattorff 2010). Hence, this implies not primarily an effect of N_e_ on substitution rates, but also other sociality-related features may impact the observed rates of molecular evolution of anti-viral siRNAi genes. Especially the high parasite pressure associated with group-living of highly related individuals, are expected to overpower the effect of N_e_ in this study, as the antiviral RNAi pathway is part of the innate immune system, where selection is expected to be particularly strong. The loss of immune genes observed for honeybees compared to *Drosophila*, is assumed to be due to alternative defense strategies such as socially mediated physiological adaptations and behavioral activities. Many of these social defense mechanisms, e.g. removal of infected individuals, collection and secretion of antimicrobial substances, serve to avoid bacterial or fungal infection (Cremer et al. 2007). Hence, features associated with social immunity may particularly reduce the selection coefficient acting on genes involved in antibacterial or antifungal defense mechanisms, but not on antiviral genes. With regard to their short “generation times” and high mutation rates viruses *per se* pose a particular challenge to host immune system. And, as honeybee viruses have been shown to infect bumblebees, there is increasing evidence for cross-species transmission and establishment (Genersch et al. 2006; Singh et al. 2010; Levitt et al. 2013; Fürst et al. 2014), which likewise might reinforce a selective pressure. For honeybees silencing of viral RNA after ingestion of viral dsRNA leads to a reduction in viral load and bee mortality (Maori et al. 2009; Desai et al. 2012) - indicating the functional importance of an effective RNAi pathway in managing viral infections. Moreover, within-species comparisons revealed only scarce evidence for polymorphisms, most of them being synonymous (Table S1), indicating rather selection than relaxed selective constraint may impact the molecular evolution of the anti-viral siRNAi genes.

Also reproductive features might contribute to the different substitution rates. In social Hymenoptera, the delayed production of sexuals is assumed to increase the chance for inheriting mutations due to accumulation of DNA copy errors, as oogenesis in Hymenoptera is continuous, which is assumed to increase mutation rates as the number of germline cell divisions increases with generation (Büning 1994; Crow 1997; Bromham and Leys 2005). In contrast, socially parasitic cuckoo bumblebees lack a worker caste and therefore, the female immediately starts with the production of reproductive offspring. Thus, the effect of continuous oogenesis on the mutation rate might not be that strong.

### Conclusion

High infection risk is one of the main challenges social insects have to counteract. In that regard their downscaled immune gene repertoire appears to be counterintuitive. Here, behavioral defense mechanisms might provide valuable support reducing infection risk and in that regard selective pressure on immune genes.

For antiviral defense mechanisms, where selection pressure is exceptionally strong and social immunity is unlikely to work efficiently, sociality associated features (high infection risk; centralized reproductive privilege causing low N_e_; consequences of continuous oogenesis and delayed sexuals production on mutation rate) seems to affect the rate of molecular evolution. Evidence is given as for three out of six siRNAi genes higher evolutionary rates were observed for the social species compared to their non-social parasitic cuckoo bumblebees. The latter ones are assumed to show higher rates due to a relaxed selective constraint as a consequence of a stronger reduction in N_e_.

## Acknowledgements

We are grateful to D. Kleber and P. Leibe for assistance with lab work, especially for NGS sequencing. We would like to thank S. Erler for providing genomic DNA of bumblebee species, which were collected by P. Lhomme and P. Rasmont. We also thank B. Fouks, P. Thoisy, B. Quaye and A. Weiss for providing sequences of intraspecific r2d2 variability. We thank the Bumblebee Genome Consortium (http://hymenopteragenome.org/beebase/) for providing genomic resources that were used for this study. This work was supported by a grant from the German Ministry for Education and Research (BMBF) within the program FUGATO-Plus (FKZ: 0315126 to HMGL).

RNAi gene sequences for all species were deposited in GenBank (accession numbers:)

## References

Barribeau, SM. et al. A depauperate immune repertoire precedes evolution of sociality in bees. Genome Biol (in press) (2014).

Boomsma JJ, Schmid-Hempel P, Hughes WOH.2005. Life Histories and Parasite Pressure Across the Major Groups of Social Insects. In Fellowes M, Holloway G, Rolff J, editors, Insect Evolutionary Ecology. Wallingford Oxon, UK & Cambridge MA, USA: CABI Publishing. p. 139–175.

Bromham L, Leys R. 2005. Sociality and the rate of molecular evolution. Mol. Biol. Evol. 22: 1393–1402. doi: 10.1093/molbev/msi133

Buchon N, Vaury C. 2006. RNAi: a defensive RNA-silencing against viruses and transposable elements. Heredity 96: 195–202. doi:10.1038/sj.hdy.6800789

Bulmer MS, Crozier RH. 2004. Duplication and diversifying selection among termite antifungal peptides. Mol. Biol. Evol. 21: 2256–2264. doi: 10.1093/molbev/msh236

Bulmer MS, Crozier RH. 2006. Variation in positive selection in termite GNBPs and Relish. Mol. Biol. Evol. 23: 317–326. doi: 10.1093/molbev/msj037

Büning J. 1994. The Insect Ovary: Ultrastructure, Previtellogenic Growth and Evolution. Springer.

Cameron SA, Hines HM, Williams PH. 2007. A comprehensive phylogeny of the bumble bees (Bombus). Biol. J. Linn. Soc. 91: 161–188. doi: 10.1111/j.1095- 8312.2007.00784.x

Caudy AA, Myers M, Hannon GJ, Hammond SM. 2002. Fragile X-related protein and VIG associate with the RNA interference machinery. Genes Dev. 16: 2491–2496. doi: 10.1101/gad.1025202

Cremer S, Armitage SA, Schmid-Hempel P. 2007. Social immunity. Curr Biol. 17: 693–702. doi:10.1016/j.cub.2007.06.008

Crow JF. 1997. The high spontaneous mutation rate: is it a health risk? Proc. Natl. Acad. Sci. USA. 94: 8380–8386.

Crozier RH. 1977. Evolutionary genetics of the Hymenoptera. AnnuRev Entomol. 22:263–288. doi: 10.1146/annurev.en.22.010177.001403

Desai S D, Eu Y-J, Whyard S, Currie RW. 2012. Reduction in deformed wing virus infection in larval and adult honey bees (Apis mellifera L.) by double-stranded RNA ingestion. Insect Mol. Biol. 21: 446–455. doi: 10.1111/j.1365-2583.2012.01150.x.

Ding S-W. 2010. RNA-based antiviral immunity. Nat. Rev. Immunol. 10: 632–644. doi: 10.1038/nri2824.

Erler S, Lattorff HMG. 2010. The degree of parasitism of the bumblebee (Bombus terrestris) by cuckoo bumblebees (Bombus (Psithyrus) vestalis). Insectes Sociaux 57: 371–377. doi: 10.1007/s00040-010-0093-2.

Erler S, Lhomme P, Rasmont P, Lattorff HMG. 2014. Rapid evolution of antimicrobial peptide genes in an insect host-social parasite system. Infect. Genet. Evol. 23: 129–137. doi: 10.1016/j.meegid.2014.02.002.

Evans JD, Aronstein K, Chen YP, Hetru C, Imler J-L, Jiang H, Kanost M, Thompson GJ, Zou Z, Hultmark D. 2006. Immune pathways and defence mechanisms in honey bees Apis mellifera. Insect Mol. Biol. 15: 645–656. doi: 10.1111/j.1365-2583.2006.00682.x.

Felsenstein J. 2005. PHYLIP (Phylogeny Inference Package) version 3.6. Distributed by the author. Department of Genome Sciences, University of Washington, Seattle.

Fürst MA, McMahon DP, Osborne JL, Paxton RJ, Brown MJF. 2014. Disease associations between honeybees and bumblebees as a threat to wild pollinators. Nature 506: 364–366. doi: 10.1038/nature12977.

Genersch E, Yue C, Fries I, de Miranda JR. 2006. Detection of Deformed wing virus, a honey bee viral pathogen, in bumble bees (Bombus terrestris and Bombus pascuorum) with wing deformities. J. Invertebr. Pathol. 91: 61–63. doi:10.1016/j.jip.2005.10.002.

Goulson D. 2010. Bumblebees: Behaviour, Ecology and Conservation. Oxford University Press.

Gracheva E, Dus M, Elgin SCR. 2009. Drosophila RISC component VIG and its homolog Vig2 impact heterochromatin formation. PLoS ONE 4: e6182. DOI: 10.1371/journal.pone.0006182.

Hall TA. 1999. BioEdit: a user-friendly biological sequence alignment editor and analysis program for Windows 95/98/NT. Nucl. Acids. Symp. Ser. 41: 95–98.

Hamilton WD. 1964. Genetical evolution of social behaviour. J. Theor. Biol. 7: 1–52.

Harpur BA, Zayed A. 2013. Accelerated evolution of innate immunity proteins in social insects: adaptive evolution or relaxed constraint? Mol. Biol. Evol. 30: 1665–1674. doi: 10.1093/molbev/mst061.

Jukes TH, Cantor CR. 1969. Evolution of protein molecules. In: Munro HN, editors. Mammalian Protein Metabolism. New York: Academic Press. p. 21–132.

Kolaczkowski B, Hupalo D N, Kern AD. 2011. Recurrent adaptation in RNA interference genes across the Drosophila phylogeny. Mol. Biol. Evol. 28: 1033–1042. doi: 10.1093/molbev/msq284.

Lattorff HMG, Moritz RFA. 2013. Genetic underpinnings of division of labor in the honeybee (Apis mellifera). Trends Genet. 29: 641–648. doi: 10.1016/j.tig.2013.08.002.

Levitt AL, Singh R, Cox-Foster DL, Rajotte E, Hoover K, Ostiguy N, Holmes EC. 2013. Cross-species transmission of honey bee viruses in associated arthropods. Virus Res. 176: 232–240. doi: 10.1016/j.virusres.2013.06.013.

Li H-W, Ding S-W. 2005. Antiviral silencing in animals. FEBS Lett. 579: 5965–5973. doi: 10.1016/j.febslet.2005.08.034.

Li F, Ding S-W. 2006. Virus counterdefense: diverse strategies for evading the RNA-silencing immunity. Annu. Rev. Microbiol. 60: 503–531. doi: 10.1146/annurev.micro.60.080805.142205.

Librado P, Rozas J. 2009. DnaSP v5: A software for comprehensive analysis of DNA polymorphism data. Bioinformatics 25: 1451–1452. doi: 10.1093/bioinformatics/btp187.

Liu Q, Rand TA, Kalidas S, Du F, Kim HE, Smith DP, Wang X. 2003. R2D2, a bridge between the initiation and effector steps of the Drosophila RNAi pathway. Science 301: 1921–1925. doi: 10.1126/science.1088710.

Maori E, Paldi N, Shafir S, Kalev H, Tsur E, Glick E, Sela I. 2009. IAPV, a bee-affecting virus associated with Colony Collapse Disorder can be silenced by dsRNA ingestion. Insect Mol. Biol. 18: 55–60. doi: 10.1111/j.1365-2583.2009.00847.x.

Meister G, Tuschl T. 2004. Mechanisms of gene silencing by double-stranded RNA. Nature 431: 343–349. doi:10.1038/nature02873.

Moissiard G, Voinnet O. 2004. Viral suppression of RNA silencing in plants. Mol. Plant Pathol. 5: 71–82. DOI: 10.1111/j.1364-3703.2004.00207.x.

Nayak A, Berry B, Tassetto M, Kunitomi M, Acevedo A, Deng C, Krutchinsky A, Gross J, Antoniewski C, Andino R. 2010. Cricket paralysis virus antagonizes Argonaute 2 to modulate antiviral defense in Drosophila. Nat. Struct. Mol. Biol. 17: 547–554. doi: 10.1038/nsmb.1810.

Nayak A, Tassetto M, Kunitomi M, Andino R. 2013. RNA interference-mediated intrinsic antiviral immunity in invertebrates. In: Cullen BR, editor. Intrinsic Immunity. Current topics in Microbiology and Immunology. 371: 183–200.

Obbard DJ, Jiggins FM, Halligan DL, Little TJ. 2006. Natural selection drives extremely rapid evolution in antiviral RNAi genes. Curr. Biol. 16: 580–585. doi:10.1016/j.cub.2006.01.065.

Obbard DJ, Gordon K H J, Buck A H, Jiggins FM. 2009a. The evolution of RNAi as a defence against viruses and transposable elements. Philos. Trans. R. Soc. Lond. B. Biol. Sci. 364: 99–115. doi: 10.1098/rstb.2008.0168.

Obbard DJ, Welch JJ, Kim KW, Jiggins FM. 2009b. Quantifying adaptive evolution in the Drosophila immune system. PLoS Genetics 5: e1000698. doi: 10.1371/journal.pgen.1000698

Ohta T. 1987. Very slightly deleterious mutations and the molecular clock. J Mol. Evol. 26: 1–6.

Romiguier J, Lourenco J, Gayral P, Faivre N, Weinert LA, Ravel S, Ballenghien M, Cahais V, Bernard A, Loire E, et al. 2014. Population genomics of eusocial insects: the costs of a vertebrate-like effective population size. J. Evol. Biol. 27: 593–603 (2014). doi: 10.1111/jeb.12331

Rozen S, Skaletsky H. 2000. Primer3 on the WWW for general users and for biologist programmers. Methods Mol Biol. 132: 365–386.

Schmid-Hempel P. 1998. Parasites in Social Insects. Princeton: Princeton University Press.

Singh R, Levitt AL, Rajotte EG, Holmes EC, Ostiguy N, vanEngelsdorp D, Lipkin WI, Depamphilis CW, Toth AL, Cox-Foster DL. 2010. RNA viruses in hymenopteran pollinators: evidence of inter-Taxa virus transmission via pollen and potential impact on non-Apis hymenopteran species. PLoS ONE 5, e14357. doi: 10.1371/journal.pone.0014357.

Smith CR, Smith CD, Robertson HM, Helmkampf M, Zimin A, Yandell M, Holt C, Hu H, Abouheif E, Benton R, et al. 2011. Draft genome of the red harvester ant Pogonomyrmex barbatus. Proc. Natl. Acad. Sci. USA 108: 5667–5672. doi: 10.1073/pnas.1007901108.

Tamura K, Peterson D, Peterson N, Stecher G, Nei M, Kumar S. 2011. MEGA5: Molecular evolutionary genetics analysis using maximum likelihood, evolutionary distance, and maximum parsimony methods. Mol. Biol. Evol. 28: 2731–2739. doi: 10.1093/molbev/msr121.

Thompson JD, Higgins DG, Gibson TJ. 1994. CLUSTAL W: improving the sensitivity of progressive multiple sequence alignment through sequence weighting, position specific gap penalties and weight matrix choice. Nucleic Acids Res. 22: 4673–4680.

van Mierlo JT, Bronkhorst AW, Overheul GJ, Sadanandan SA, Ekström JO, Heestermans M, Hultmark D, Antoniewski C, van Rij RP. 2012. Convergent evolution of argonaute-2 slicer antagonism in two distinct insect RNA viruses. PLoS Pathog. 8: e1002872. doi: 10.1371/journal.ppat.1002872.

Viljakainen L, Evans JD, Hasselmann M, Rueppell O, Tingek S, Pamilo P. 2009. Rapid evolution of immune proteins in social insects. Mol. Biol. Evol. 26: 1791–1801. doi: 10.1093/molbev/msp086.

Wilson EO. 1990. Success and dominance in ecosystems: the case of the social insects. Oldendorf/Luhe.

Yang Z. 2007. PAML 4: a program package for phylogenetic analysis by maximum likelihood. Mol. Biol. Evol. 24: 1586–1591. doi: 10.1093/molbev/msm088.

Yang Z, Kumar S, Nei M. 1995. A new Method of inference of ancestral nucleotide and amino acid sequences. Genetics 141: 1641–1650.

Zhang J, Kumar S. 1997. Detection of convergent and parallel evolution at the amino acid sequence level. Mol. Biol. Evol. 14: 527–536.

Zhang J, Nei, M. 1997. Accuracies of ancestral amino acid sequences inferred by the parsimony, likelihood, and distance methods. J. Mol. Evol. 44: 139–146.

